# Nonadaptive radiation of the gut microbiome in an adaptive radiation of *Cyprinodon* pupfishes with minor shifts for scale-eating

**DOI:** 10.1101/2021.01.06.425529

**Authors:** J. Heras, C.H. Martin

## Abstract

Adaptive radiations offer an excellent opportunity to understand the eco-evolutionary dynamics of gut microbiota and host niche specialization. In a laboratory common garden, we compared the gut microbiota of two novel trophic specialists, a scale-eater and a molluscivore, to a set of four outgroup generalist populations from which this adaptive radiation originated. We predicted an adaptive and highly divergent microbiome composition in the specialists matching their rapid rates of craniofacial diversification in the past 10 kya. We measured gut lengths and sequenced 16S rRNA amplicons of gut microbiomes from lab-reared fish fed the same high protein diet for one month. In contrast to our predictions, gut microbiota largely reflected 5 Mya phylogenetic divergence times among generalist populations in support of phylosymbiosis. However, we did find significant enrichment of *Burkholderiaceae* bacteria in both lab-reared scale-eater populations. These bacteria sometimes digest collagen, the major component of fish scales, supporting an adaptive shift. We also found some enrichment of *Rhodobacteraceae* and *Planctomycetacia* in lab-reared molluscivore populations, but these bacteria target cellulose. Minor shifts in gut microbiota appear adaptive for scale-eating in this radiation, whereas overall microbiome composition was phylogenetically conserved. This contrasts with predictions of adaptive radiation theory and observations of rapid diversification in all other trophic traits in these hosts, including craniofacial morphology, foraging behavior, aggression, and gene expression, suggesting that microbiome divergence proceeds as a nonadaptive radiation.

## Introduction

Rapid evolutionary change can alter ecological processes which in turn change the course of evolutionary processes (Turcotte et al. 2013; Matthews et al. 2016). This process is described as eco-evolutionary dynamics, which provides a framework for understanding the interplay between evolution and ecological interactions (Rudman et al. 2018; Post et al. 2009). The emergence of studies that focus on eco-evolutionary dynamics has provided more insight on the processes of community assembly, ecological speciation, and adaptive radiations (Rudman et al. 2018). A better understanding of these eco-evolutionary dynamics can be applied to host-microbiota interactions, in which the co-evolutionary processes of the microbiome can impact host performance and fitness (Gould et al. 2018; Macke et al. 2017; Walters et al. 2020). The microbial community may also play a large role in the physiology, ecology, and evolution of the host (Baldo et al. 2017; Trevelline and Kohl 2020).

Several studies have now examined gut microbiome diversification in an adaptive radiation of hosts, including fishes (Baldo et al. 2017; Baldo et al. 2019; Loo et al. 2019; Macke et al. 2017; Rennison et al. 2019). Phylosymbiosis, in which the host microbiome recapitulates host phylogeny, is frequently the primary hypothesis in these studies (Brooks et al. 2016; Lim and Bordenstein, 2020). However, these studies rarely examine outgroups to the focal radiation in order to compare rates of microbiome divergence. Furthermore, phylosymbiosis (comparable to phylogenetic conservatism; Losos, 2008) is actually the antithesis to the theory of adaptive radiation, which predicts that the microbiome within an adaptive radiation should diverge far more quickly than outgroup taxa due to rapid ecological divergence and specialization (Stroud and Losos 2016, Schluter 2000, Martin and Richards 2019, Gillespie et al. 2020, Rundell and Price 2009). Thus, we predicted greater microbiome divergence within a recent adaptive radiation of trophic specialists than in outgroup generalist taxa with far older divergence times (5 Mya), in contrast to the predictions of phylosymbiosis.

An adaptive radiation of *Cyprinodon* pupfishes provides an excellent opportunity to test the relative roles of rapid trophic divergence and phylosymbiosis in shaping the gut microbiome. Pupfishes are found in saline lakes or coastal areas throughout the Caribbean and Atlantic (most are allopatric) and within isolated desert pools and streams (Martin et al. 2016, 2020; Echelle and Echelle 2020). However, there are only two sympatric adaptive radiations of trophic specialists across this range (Martin and Wainwright 2011). One radiation is endemic to San Salvador Island, Bahamas, containing a generalist algivorous and detritivorous species, *Cyprinodon variegatus,* and two trophic specialist species, a molluscivore *C. brontotheroides* and a scale-eater *C. desquamator* (Martin and Wainwright, 2011; Martin and Wainwright, 2013; Richards and Martin, 2017). Scale-eating and molluscivore niches are uniquely derived within this sympatric radiation on San Salvador Island relative to generalist outgroup populations spread across the Caribbean and desert interior of North America (Martin and Feinstein 2014; Richards and Martin 2016). These two specialist species diverged from a generalist common ancestor within the past 10 kya, drawing adaptive alleles from ancient standing genetic variation across the Caribbean (Richards et al. 2020; McGirr and Martin 2020), whereas the most divergent generalist population in our study, the checkered pupfish *Cualac tessellatus,* has persisted for up to 5 Mya in El Potosí desert spring system in Mexico (Echelle et al. 2005). Thus, this radiation provides an excellent opportunity to compare microbiome divergence within a sympatric adaptive radiation of trophic specialists to closely related and ancient outgroup generalist taxa which have not substantially shifted their dietary niches.

We compared gut length, overall microbiome diversity, and enrichment of specific microbial taxa among three sympatric *Cyprinodon* pupfish species from two different isolated lake populations on San Salvador Island, Bahamas to three generalist species: closely related *C. laciniatus* from Lake Cunningham, New Providence Island, Bahamas; more distantly related C. *variegatus* from Fort Fisher, North Carolina; and the most closely related extant genus *Cualac tessellatus* from San Luis Potosí, Mexico. We raised all these species in a common laboratory environment for at least one generation and fed them an identical commercial pellet diet for one month before sampling gut microbiomes. We addressed the following questions: 1) Do microbial gut communities vary by diet or phylogenetic distance among these species? 2) Is there a microbiome signal associated with lepidophagy (scale-eating) or molluscivory?

## Materials and Methods

### Sampling and preparation of gut microbiome samples

Colonies of *Cyprinodon* pupfishes were collected from two hypersaline lakes on San Salvador Island, Bahamas (Crescent Pond and Osprey Lake) and Lake Cunningham, Bahamas in March, 2018 and were reared in aquaria at the University of California, Berkeley. Additional generalist populations were collected in May, 2018 from Fort Fisher Estuary in North Carolina. *Cualac tessellatus* eggs were provided by the Zoological Society of London and reared in the lab to produce a large second generation used for the four samples in this study. All samples, except for the recently collected NC population, came from first or second-generation captive-bred individuals reared in aquaria (40–80 L) according to species and location at 5–10 ppt salinity (Instant Ocean synthetic sea salt) and between 23 to 30°C. Individuals used for this study were first fed once daily *ad libitum* with a single commercial pellet food (New Life Spectrum Cichlid Formula, New Life International, Inc., Homestead, FL), containing 34% crude protein, 5% crude fat, and 5% crude fiber, for one month without exposure to any other food or tankmates. All animal care and experiments were conducted under approved protocols and guidelines of the University of California, Berkeley Institutional Animal Care and Use Committee (AUP-2018-08-11373).

In total, forty fishes were euthanized in an overdose of MS-222 and the entire intestinal tissue was immediately excised (Cyprinodontidae do not possess stomachs; Wilson and Castro, 2010) for DNA extraction. Standard length and gut length were measured for all samples. Five individuals (F_2_ generation) from each of three species (*C. variegatus, C. brontotheroides,* and *C. desquamator)* in both lake populations from San Salvador Island were sampled (*n* = 30 total). In addition, we included the following pupfish species as outgroups to our study: *C. laciniatus* (F1 generation; Lake Cunningham, New Providence Island, Bahamas; *n* = 4), *C. variegatus* (F0 generation; Fort Fisher, North Carolina, United States; *n* = 2) plus liver tissue as a tissue control, and *Cualac tessellatus* (long-term captive colony; San Luis Potosí, Mexico, *n* = 4).

Each gut was divided into proximal and distal regions for all San Salvador Island samples to compare microbial composition between these regions. All outgroup samples used whole intestines. In addition, the microbial community was isolated from aquaria water in two tanks which contained F2 individuals of Osprey Lake *C. variegatus* and Crescent Pond *C. variegatus,* and used as controls (*n* = 2). The Vincent J. Coates Genomics Sequencing Laboratory at the University of California, Berkeley also generated three controls, including a positive control and two no template controls (NTC). Microbial DNA extractions were performed in batches (stored on ice) immediately after intestinal dissections with the Zymobiomics DNA Miniprep Kit (Zymo Research, Irvine, CA).

### 16S amplicon sequencing of gut microbiomes

All extracted microbiome DNA samples were quantified with a Nanodrop ND-1000 spectrophotometer (range 4.2-474.9 ng/μl). All samples were then sent to the QB3 Vincent J. Coates Genomics Sequencing Laboratory at the University of California, Berkeley for automated library preparation and sequencing of 16S rRNA amplicons using an Illumina Mi-Seq v3 (600 cycle). As part of the QB3 library preparation, the Forward ITS1 (ITS1f) – CTTGGTCATTTAGAGGAAGTAA and Reverse ITS1 (ITS2) – GCTGGGTTCTTCATCGATGC primers (Smith and Peay, 2014) were used for DNA metabarcoding markers for fungi (Smith and Peay, 2014). QB3 also used the following 16S rRNA primers for amplification of prokaryotes (archaea and bacteria): Forward 16S v4 (515Fb) – GTGYCAGCMGCCGCGGTAA, and Reverse 16S v4 (806Rb) – GGACTACNVGGGTWTCTAAT (Caporaso et al., 2011; Apprill et al. 2015).

### Bioinformatic Analysis/Quantification and Microbial Ecology Assessment of Samples

All 16S rRNA amplicon sequences were processed through QIIME 2.0 (Bolyen et al. 2018) to identify microbe species and estimate abundances. Sequences from all 78 microbiome preps were imported into QIIME v. 2019.10.0. We determined there were no differences between proximal and distal regions of the gut for the San Salvador Island individuals, therefore we concatenated the Crescent Pond and Osprey Lake samples into one file, in which we had 48 samples which included experimental controls and quality controls from the QB3 facility (Table S2). There was no difference between the means of microbe counts in the foregut and the hindgut (paired t-test, *P* = 0.29).

We used DADA2 (Callahan et al. 2016) for modeling and correcting Illumina-sequenced amplicon errors, removing chimeras, trimming low quality bases, and merging of forward and reverse reads using the following parameters: −p-trunc-len-f 270 −p-trunc-len-r 210. We used the QIIME *alignment mafft* software to align sequences *alignment mask* to filter non-conserved and highly gapped columns from the aligned 16S sequences (Stackebrandt and Goodfellow, 1991). Next, we used *qiime phylogeny midpoint-root* to root the phylogeny of our 16S amplicon sequences. Finally, we used *qiime diversity alpha-rarefaction* on all samples and we set the --p- max-depth to 10,000. We removed samples with 5,000 or less from our analyses.

We compared the beta diversity (*qiime emperor plot*) of proximal and distal gut microbiomes of the San Salvador samples with a two-tailed paired *t*-test and found no significant differences between proximal and distal regions of the gut microbiome (*P* = 0.29). Therefore, we merged the proximal and distal samples for each individual from San Salvador Island, resulting in 48 samples. We also removed one *Cualac tessellatus* sample because of low read count (129 reads; Figure S2).

We used the classifier Silva 132 99% 515F/806R (silva-132-99-515-806-nb-classifier) for training in identification of taxa from our samples. Afterwards the following files generated in QIIME were used in R (v. 4.0.0) for further statistical analyses: table.qza, rooted-tree.qza, taxonomy.qza, and sample-metadata.tsv. We used the following R packages for further analyses: phyloseq (McMurdie and Holmes 2013) and ggplot2 (Wickham, 2016) with the following functions: *distance, plot bar, plot ordination,* and *plot richness.* Before conducting any analyses, we removed the following taxa from our analyses, uncharacterized and Opisthokonta (eukaryotic sequences mainly due to fish 16S amplicons). We estimated alpha diversity by using the *plot richness* function and the Chao1 and Shannon’s diversity indices. For beta diversity, we used the *plot ordination* function and non-metric multidimensional scaling (NMDS) based on Bray-Curtis distances among samples. Hierarchical clustering was generated with the distance function along with *hclust* as part of fastcluster (Müllner, 2013) using the average linkage clustering method. The *plot bar* function in the phyloseq package was used to visualize relative abundance of taxa. In our taxa plots we removed abundance counts of less than 400 from our analyses. We used *ggplot2* to generate all figures (Wickham, 2016). Lastly, we used the Linear discriminant analysis Effect Size (LEfSe version 1.0; Segata et al. 2011) algorithm to identify microbial taxa that were significantly enriched in each of our specialists (scale-eater and molluscivore) in comparison to all other samples. This analysis was used to determine the features (i.e. organisms, clades, operational taxonomic units) to explain differences in assigned metadata categories. We used the nonparametric factorial Kruskal-Wallis rank-sum test to detect taxa with significant differential abundances between specialist samples and all generalist samples (scale-eaters versus generalists + molluscivores, molluscivores versus generalists + scale-eaters). We then used a Wilcoxon test for all pairwise comparisons between taxa within each significantly enriched class to compare to the class level. From the standard and gut length measurements, we used ANCOVA in R (v. 4.0.0) to test whether there was a significant difference among species based on gut length and standard length.

Lastly, we used generalized linear models (GLMs) in R to test the effects of diet (generalist, scale-eater, molluscivore), the fixed effect of location (Osprey Lake, San Salvador Island; Crescent Pond, San Salvador Island; Lake Cunningham, New Providence Island; Fort Fisher, North Carolina; and San Luis Potosí, Mexico), and their interaction on the response variables of principal coordinates axes 1 and 2.

## Results

### Intestinal lengths among species did not vary

There was no significant difference in gut lengths among the species sampled (Fig. S1; ANCOVA with covariate of log-transformed SL; *F_533_* = 0.916, *P* = 0.483).

### Gut microbiome diversity and divergence among taxa

We sequenced a total of 11,152,147 reads across all samples (Table S2). We identified 5,174 bacterial taxa in 48 samples. Similar to other ray-finned fishes (Youngblut et al. 2019), proteobacteria is the predominate microbial taxon (Figure S3). We did not find any significant differences among species in Chao1 or Shannon diversity indices (Kruskal-Wallace [pairwise], *P* > 0.05). San Salvador Island pupfishes clustered together relative to the three outgroup generalist species, indicating strong host phylogenetic signal associated with overall microbiome diversity (Fig. 2). Water and tissue controls were scattered throughout the NMDS plots but were clearly distinct from *Cyprinodon* microbiome samples with the exception of one tissue control that clustered near the outgroup species, possibly due to contamination during dissections (Fig. 2).

**Figure 1:**
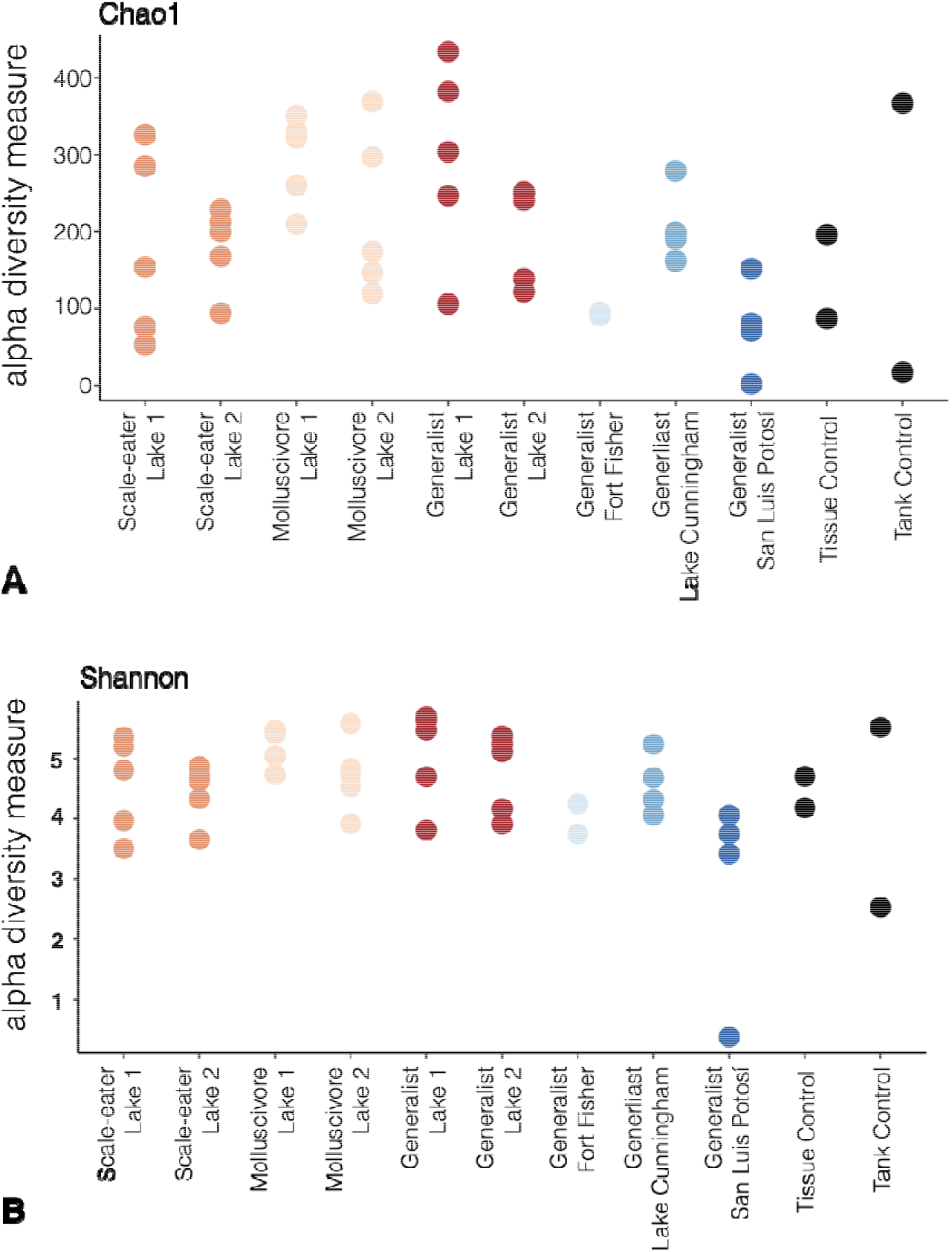
Alpha diversity of *Cyprinodon* pupfishes gut microbiomes based on parental location and diet type along with controls. Lake 1 indicates Crescent Pond and Lake 2 represents Osprey Lake, both located on San Salvador Island in the Bahamas. Alpha diversity is represented by (A) Chao1 and (B) Shannon diversity for the estimate of species richness from gut microbiomes from all fishes in this study.

**Figure 2:**
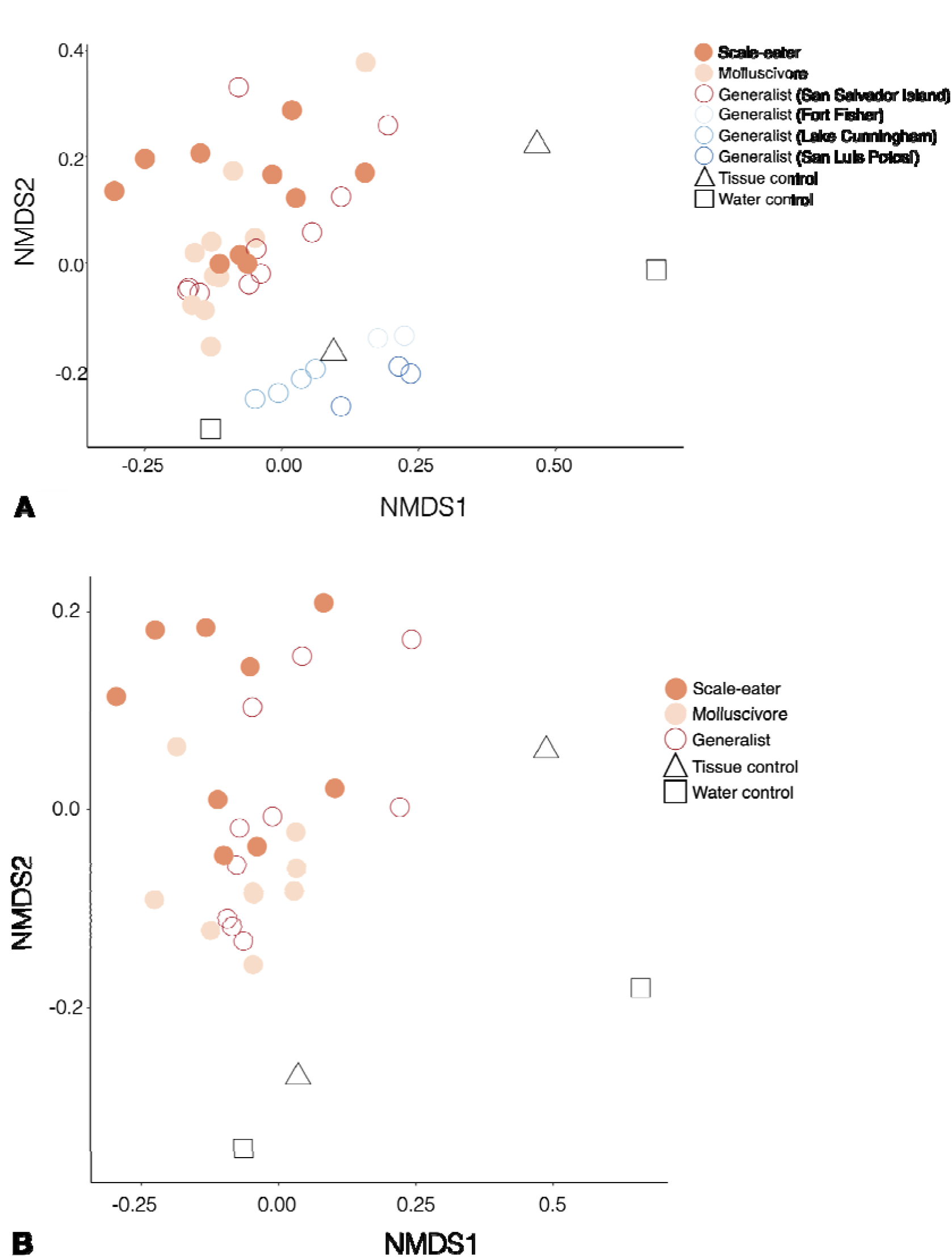
Non-metric multidimensional scaling (NMDS) plots of *Cyprinodon* pupfish gut microbiomes. A) NMDS plot based on all *Cyprinodon* pupfish gut samples labeled according to species and diet including controls (*n* = 43). B) NMDS plot of the three *Cyprinodon* pupfish species (F2 generation) from San Salvador Island including controls (*n* = 34). Closed circles represent the two specialists (scale-eater and molluscivore) and open circles represent generalists. Open squares and triangles represent controls used in this study.

Multiple regression analyses of the effects of dietary specialization (generalist, scale-eater, or molluscivore) and the fixed effect of population origin (two different lakes on San Salvador Island, Lake Cunningham, North Carolina, and El Potosí) on NMDS axes 1 and 2 confirmed that population origin and scale-eating had a significant effect on microbiome divergence along both axes (NMDS1: scale-eaters *P* = 0.001; NMDS2: scale-eaters *P* = 0.018).

### Linear discriminate analyses of trophic specialist microbiota

We found that an excess of taxa in the family *Burkholderiaceae* best discriminated all lab-reared scale-eater individuals in two different lake populations from all other gut microbiome samples (Figs. 3-4; linear discriminant analysis log score = 4.85). In addition, we found a deficiency of *Vibrionales, Vibrionaceae*, and *Vibrio* in these scale-eater individuals relative to all other gut samples (LDA log scores = −5.22, −5.22, and −5.08, respectively; Fig. 4). Similarly, we found an excess of taxa in the family *Rhodobacteraceae* and class *Planctomycetacia* in the molluscivores relative to all other gut samples (Fig. 5; LDA log scores of 4.39 and 4.37, respectively).

**Figure 3:**
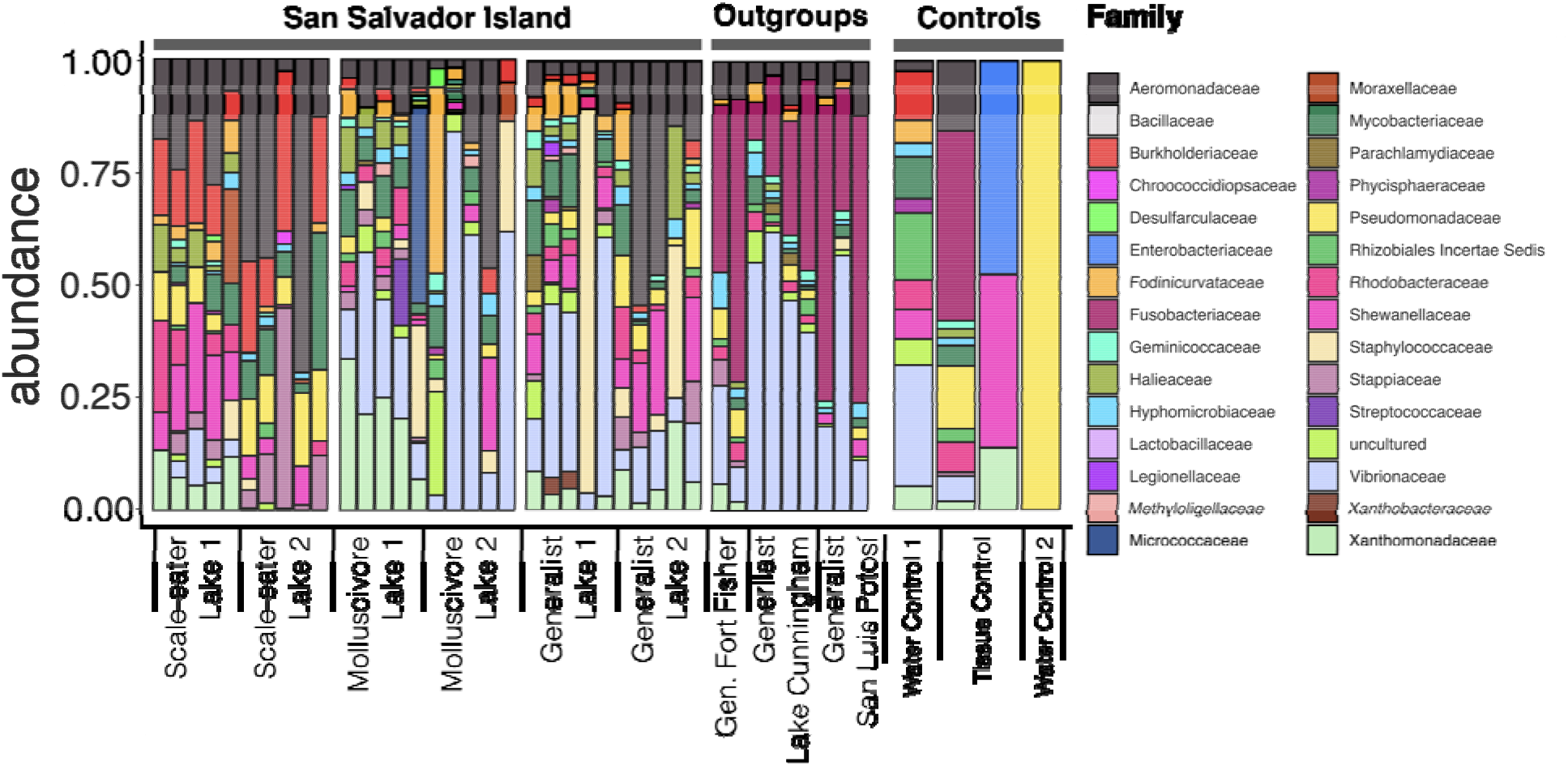
Taxa plot of the microbial composition of the *Cyprinodon* gut microbiome and controls. Bars show proportions (relative abundance) of taxa at the family level per individual gut microbiome. Lake 1 indicates Crescent Pond and Lake 2 represents Osprey Lake, both located on San Salvador Island in the Bahamas. Taxa which contained uncharacterized and Opisthokonta (eukaryotic sequences) were removed and taxa with a count of 400 or greater were represented. Taxa were grouped according to species and location (controls included).

**Figure 4:**
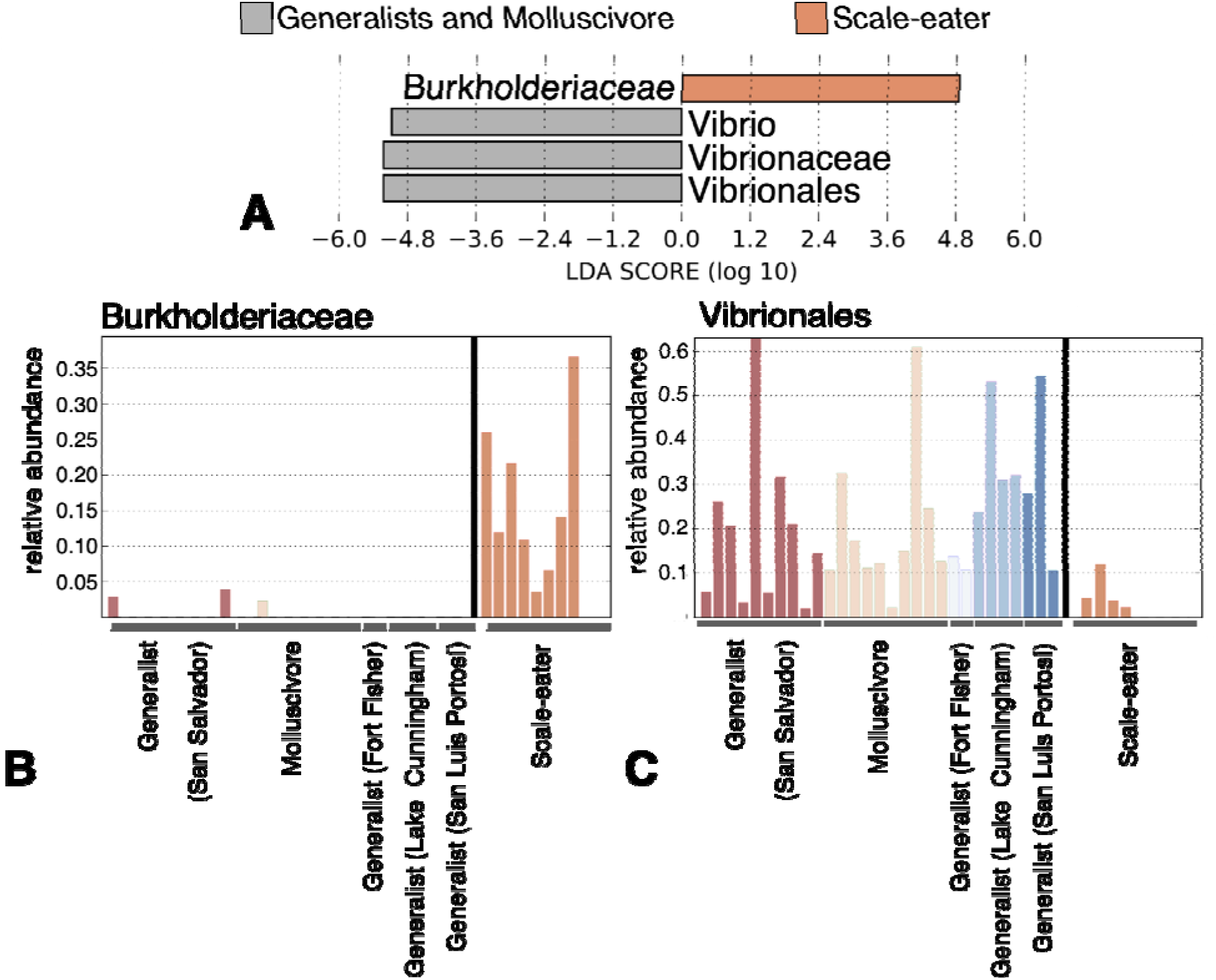
Linear discriminate analysis between *Cyprinodon desquamator* (scale-eater) and non-scale eaters. A) Log scores of the top four dominant loadings on LEfSe discriminate axis separating scale-eaters from all other pupfish samples. B) Relative abundance of the family *Burkholderiaceae* and the order C) *Vibrionales* among all pupfish gut microbiomes.

**Figure 5:**
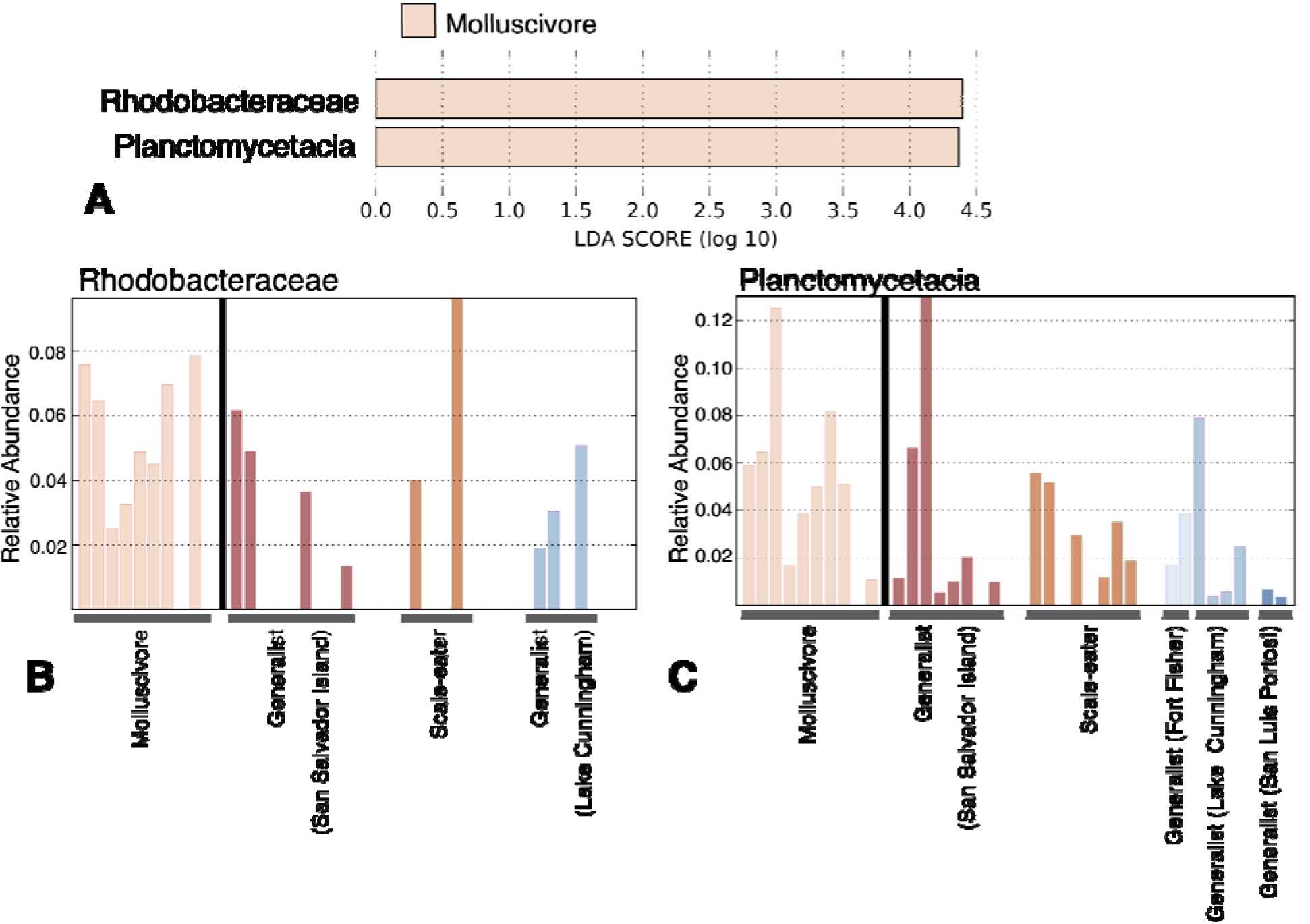
Linear discriminate analysis between *Cyprinodon brontotheroides* (molluscivore) and non-molluscivores. A) Log scores of the top two dominant loadings on the LEfSe discriminate axis separating molluscivores from all other pupfish samples. B) Relative abundance of the family *Rhodobacteraceae* and the class C) *Planctomycetacia* from all *Cyprinodon* gut microbiomes.

## Discussion

Using a common garden experiment we show that differences in gut microbial diversity across *Cyprinodon* pupfish species largely reflect phylogenetic distance among generalist populations in support of phylosymbiosis (Bordenstein and Theis 2015), rather than novel trophic specializations as predicted by adaptive radiation theory. Our study is highly consistent with Ren et al. (2016) which also found limited microbiome divergence and minimal associations with ecomorph in an adaptive radiation of Puerto Rican *Anolis* lizards, even within wild lizards. Gut microbiome diversity has also been found to associate more strongly with geography than phylogeny (Godoy-Vitorino et al., 2012) or a combination of geography, diet, and host phylogeny (Antonopoulou et al., 2019). These emerging studies of microbiome divergence within adaptive radiations of hosts provide an important counterpoint to the classic expectation of rapid phenotypic diversification and speciation during adaptive radiation (Schluter 2000; Stroud and Losos 2016; Martin and Richards 2019; Gillespie et al. 2020).

A major caveat is that we did not examine the microbiota of wild-collected animals feeding on their diverse natural resources of macroalgae, scales, and snails. Scales form up to 50% of the diet in scale-eaters (Martin and Wainwright 2013) and wild gut microbiome samples surely would have revealed more substantial differences in microbiome diversity and composition among generalist and specialist species on San Salvador Island. However, our goal with this common garden study using lab-reared animals fed an identical generalist-type diet for one month was to uncover any genetically based microbiome differences in these taxa by eliminating environmental effects as much as possible. Pupfishes exhibit no parental care and deposit external eggs on the substrate so vertical transmission also appears highly unlikely (but see Satoh et al. 2019 for a potential example of vertical transmission in a scale-eating cichlid). Furthermore, by including two lab-reared colonies of each generalist and specialist species on San Salvador from genetically differentiated and ecologically divergent lake populations (Martin et al. 2016; Richards and Martin 2017), we aimed to connect significant differences in microbiome composition observed in our specialist species to their specialized diets, rather than their lake environment or genetic background. This provides strong evidence of genetic divergence in the host associated with trophic specialization. These results are all the more surprising because trophic specialists show very little genetic differentiation from generalists (*F_st_* = 0.1 – 0.3; Martin and Feinstein 2014; Richards et al. 2020). Indeed, there are only a few thousand nearly fixed or fixed SNPs (*F_st_* > 0.95) between scale-eaters and molluscivores out of over 10 million segregating SNPs and as few as 157 fixed SNPs and 87 deletions in scale-eaters (McGirr and Martin 2020). However, this minimal set of genetic differences may be driving differences in gut microbiome composition. Intriguingly, the only fixed coding indel uncovered so far in this system is a fixed deletion in all scale-eater populations of the fifth exon of the gene gpa33 (McGirr and Martin 2020). This gene is expressed exclusively in the intestinal epithelium and mice knockouts display a range of inflammatory intestinal pathologies in mice (Williams et al. 2015), suggesting it may play a role in shifting the gut microbiota of scale-eaters that we observed in this study. Overall, metabolic processes were the single most enriched category among all differentially expressed genes between these trophic specialists at the 8 dpf larval stage, accounting for 20% of differential expression (McGirr and Martin 2018).

### Adaptive microbiota in scale-eating pupfish

Fish scales are composed of a deep layer that is mostly collagen type I (Harikrishna et al. 2017); therefore, we predicted that any adaptive microbes within the scale-eater gut would have collagen degrading properties. This includes *Bacillus, Clostridium,* and *Vibrio* taxa, which are well-known for microbial collagenase enzymes (Duarte et al. 2016). We found a significant reduction of *Vibrio* taxa within the scale-eater gut from both lake populations (Figs. 3-4). Although it is not clear why there are fewer taxa, the significant shift in a major collagenase-producing group suggests the potential for an adaptive scale-eater microbiome, even in the absence of dietary scales (except perhaps incidental aggression and ingestion of scales among tankmates). We also found significant enrichment of the family *Burkholderiaceae* in both scale-eater populations (Figs. 3-4). *Burkholderiaceae* is a family of proteobacteria which contains many human and animal pathogens (diCenzo et al. 2019), plant and insect symbionts (Gyaneshwar et al. 2011; Takeshita and Kikuchi 2017), and can be found in soil, water, and polluted environments (Coenye and Vandamme 2003; Estrada-de los Santos et al. 2016). They also include some collagenase-producing bacteria, such as *Burkholderia pseudomallei* (UniProtKB – A3P3M6; Rainbow et al. 2004), which is the causative agent of melioidosis in humans (Holden et al. 2004).

In contrast to a microbiome study of the adaptive radiation of Tanganyikan cichlids (Baldo et al. 2015), we found no evidence of *Clostridia* enrichment in scale-eaters nor a reduction of microbial diversity in this carnivorous species. This may be due to the very young 10 kya age of the scale-eating pupfish relative to the comparatively ancient 12 Mya Tanganyikan radiation and Perissodus scale-eating clade (Koblmueller et al. 2007; Martin and Wainwright 2013).

### Nonadaptive microbiota in molluscivore pupfish

We found enrichment of the families *Rhodobacteraceae* and *Planctomycetacia* within the molluscivore gut from both lake populations. However, these families have no clear role in anything related to mollusc digestion or even increased levels of protein, lipids, or chitin in the diet (due to some molluscivores specializing on ostracods during periods of abundance). Taxa from these taxonomic group are known to be found within aquatic environments (Simon et al. 2017; Yilmaz et al. 2016). Marine *Rhodobacteraceae* have a key role in biogeochemical cycling, make up about 30% of bacterial communities in the pelagic environment, and generally have a mutualistic relationship with eukaryotes providing vitamins to these groups (Simon et al. 2017). Both families are known for aquatic cellulose-decomposing taxa (Ringø et al. 2016; Kim et al. 2016), which suggests this microbiome shift may help more with macroalgae digestion rather than molluscs, despite previous observations that macroalgae forms the largest component of the generalist pupfish diet in the hypersaline lakes of San Salvador Island, Bahamas (Martin and Wainwright 2013).

### Conclusion

Many studies have focused on understanding digestion and assimilation within a variety of vertebrates and invertebrates, but there is limited information about the cooperative process between the host intestine cells and gut microbiota, and their role in eco-evolutionary dynamics during rapid species diversification (German et al. 2015; Terra et al. 2019; Baldo et al. 2017). We found evidence for a genetically-based adaptive shift in the scale-eater microbiome, even when hosts were reared in identical environments on identical non-scale diets. However, it is still unknown to what extent this microbiome shift will improve digestion of the collagen found in scales, for example, as demonstrated for the gut fauna in the scale-eating khavalchor catfish (Gosavi et al. 2018). Despite unique and highly specialized pupfish dietary adaptations within shared hypersaline lake habitats, overall gut microbial diversity did not follow the expected pattern of rapid diversification and divergence as observed in their hosts, calling into question how eco-evolutionary dynamics between host and symbiont proceed during adaptive radiation.

## Acknowledgements

This research was supported by NSF CAREER Award 1938571 and NIH/NIDCR R01 DE027052 grants to CHM. We thank L. Smith in the Evolutionary Genetics Lab at the University of California, Berkeley, for generous logistical assistance in preparing microbiome samples; S. McDevitt, C. Miller, and D. Pappas at the Vincent J. Coates Genomics Sequencing Laboratory California Institute for Quantitative Biosciences (QB3) for processing our microbiome samples for 16S amplicon sequencing; R. Berlemont at California State University, Long Beach and C. Weihe from the Microbiome Consortium at the University of California, Irvine for suggestions on microbiome extraction protocols and bioinformatic workflow. We thank the Zoological Society of London for providing *C. tessellatus* eggs and the governments of the Bahamas and United States for permission to collect and export *Cyprinodon* samples.

## Data Accessibility

Data will be deposited to Dryad and NCBI SRA.

## Author Contributions

JH prepared all samples for sequencing, conducted statistical analyses, and wrote the manuscript. CHM revised the manuscript, acquired samples, and provided funding. Both authors designed the study.

## Supplementary Table and Figures

**Supplemental Figure 1:**
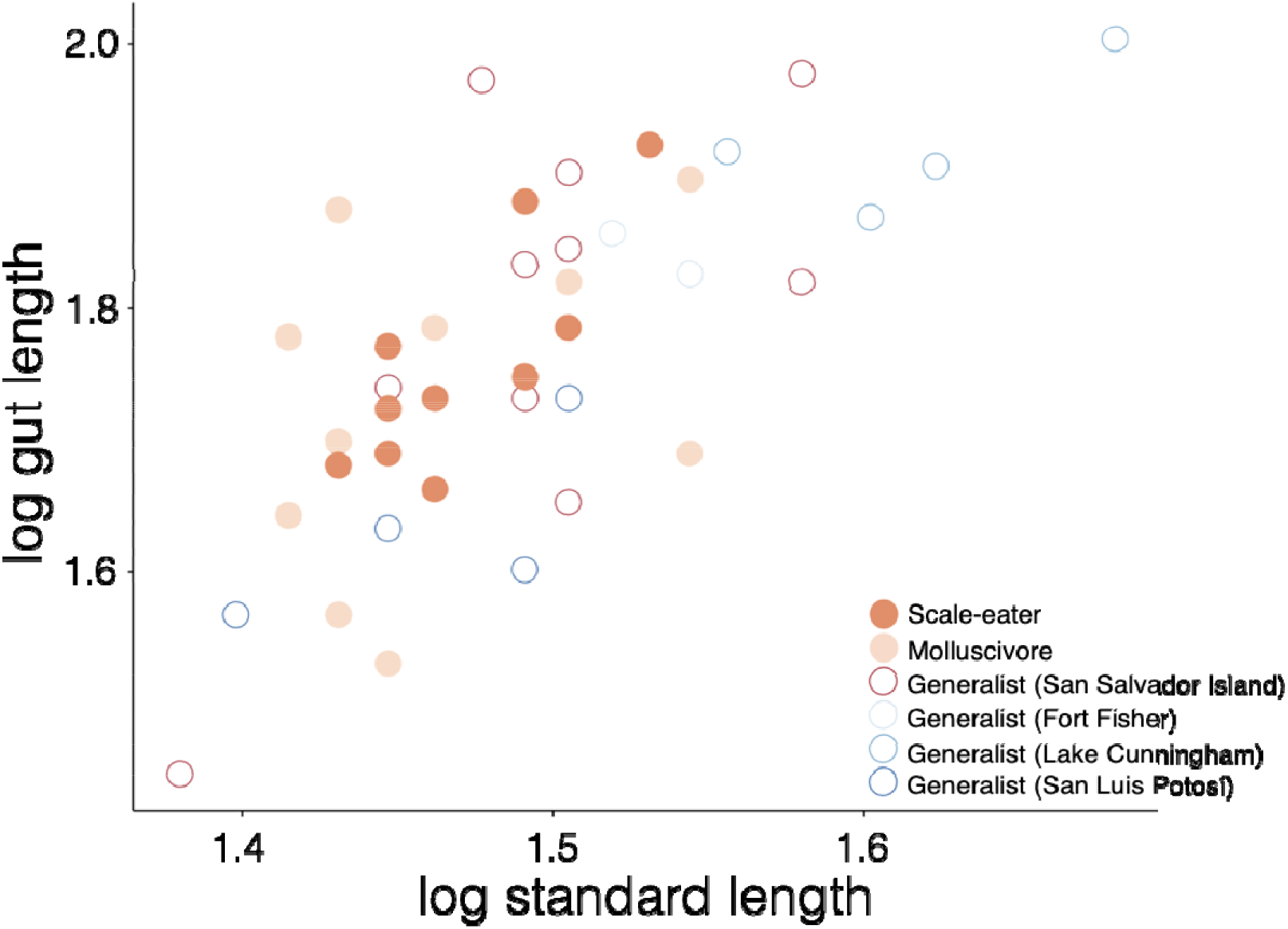
Scatter plot of the covariate (log standard length) and the outcome variable (log gut length) for all *Cyprinodon* pupfish species in our study. Closed circles represent the two specialists (scale-eater and molluscivore) and open circles represent generalists.

**Supplemental Figure 2:**
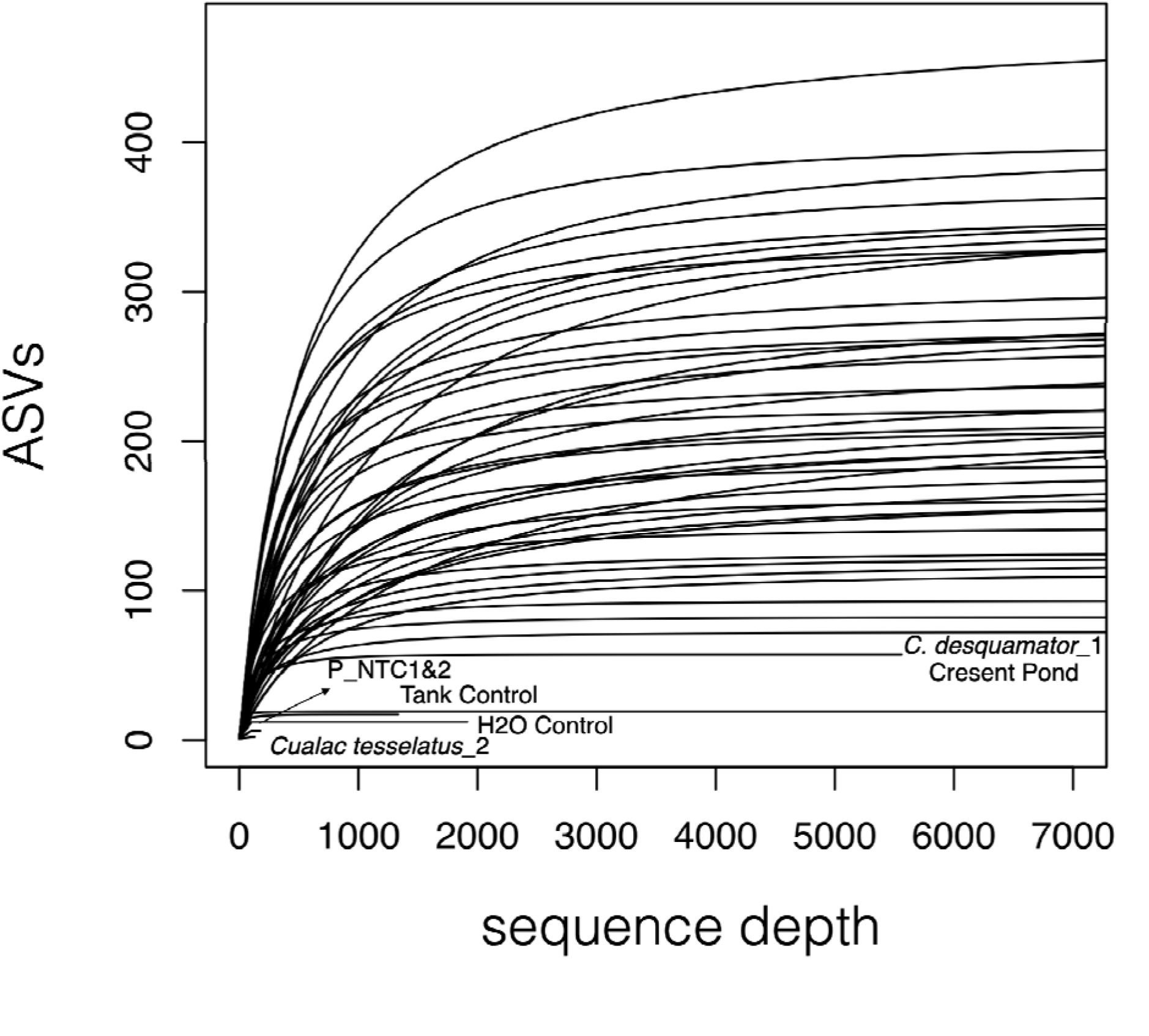
Rarefaction for all 48 (16S) microbiome samples used in this study. Rarefaction curve constructed based on amplicon Sequence Variant (ASVs), and samples with less than 6,000 reads (sequence depth) are shown with labels.

**Supplemental Figure 3:**
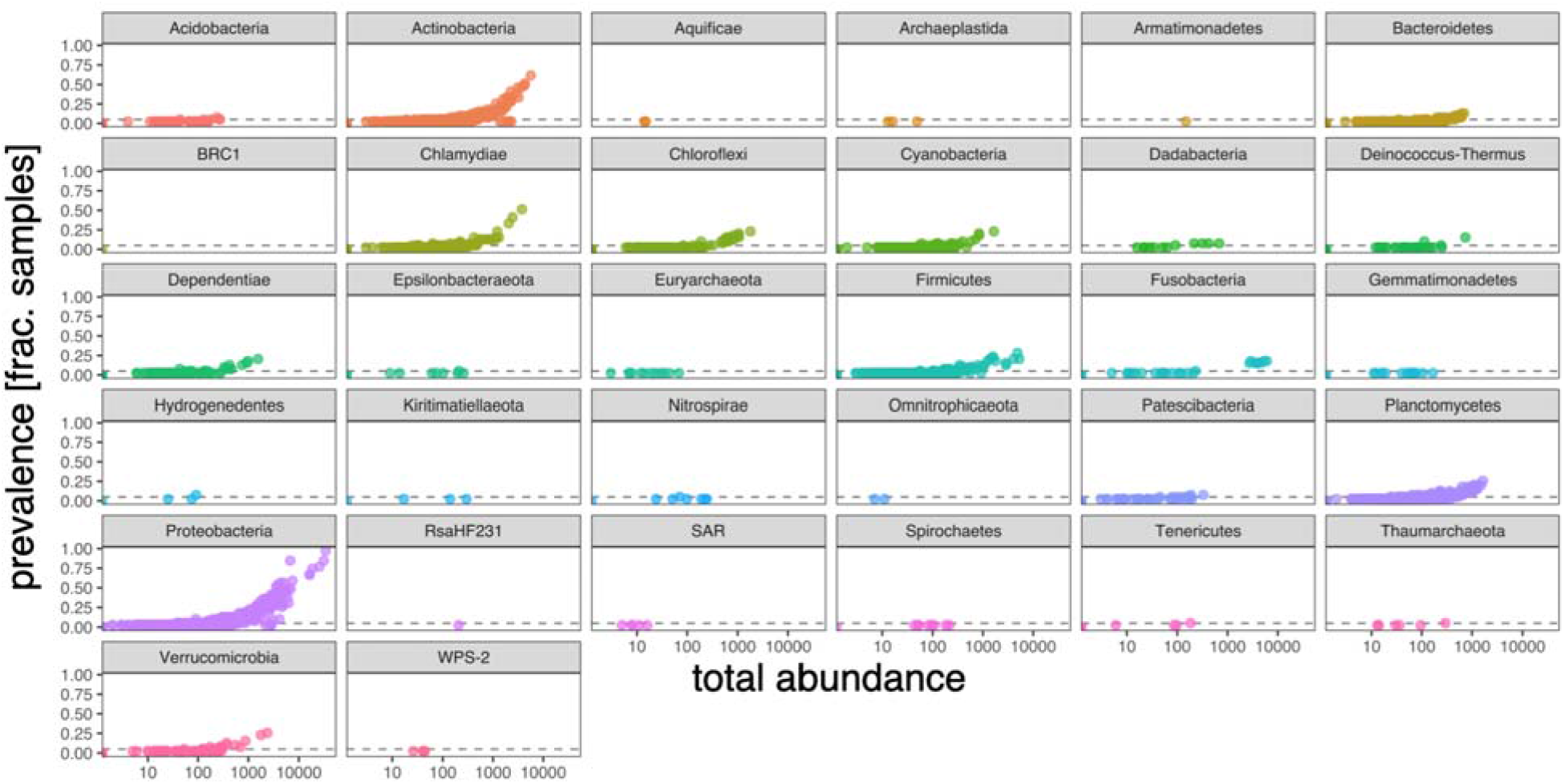
Total abundance of gut microbes across all *Cyprinodon* pupfish species used in this study. Thirty-two phyla of microbes represented across all gut microbiomes, not including controls.

**Supplemental Figure 4:**
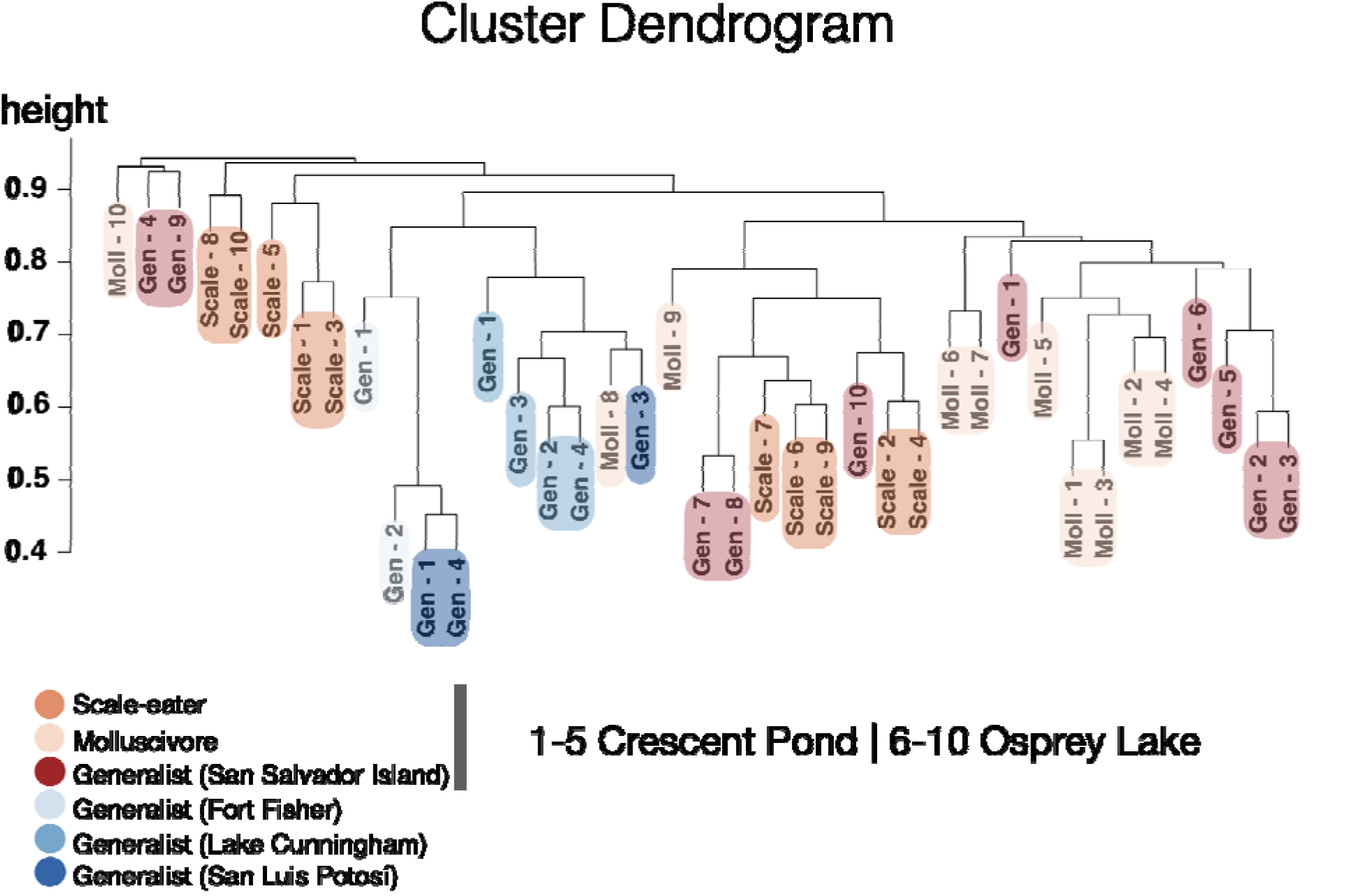
A cluster dendrogram based on pupfish gut microbiome taxa using a Bray-Curtis distance (averaged). For the San Salvador Island samples only, individuals numbered as 1-5 represent Crescent Pond and 6-10 represent Osprey Lake. Scale = scale-eater, Moll = molluscivore, and Gen = generalist.

**Supplemental Table S1:**
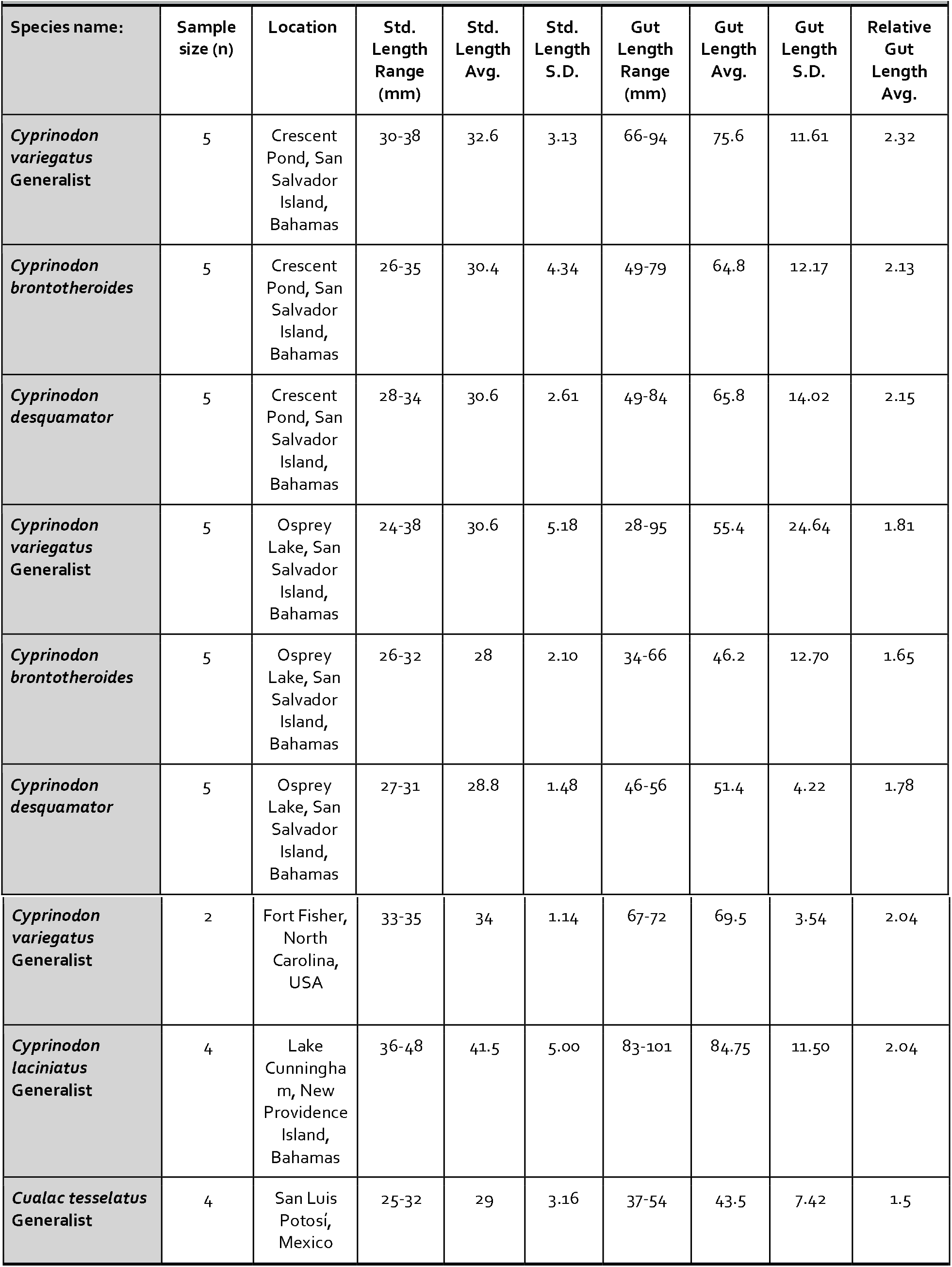
Sample Size, Location, Standard Length, Gut Length, and Relative Gut Length of *Cyprinodon* pupfish guts

|  |  |  |  |  |  |  |  |  |  |
| --- | --- | --- | --- | --- | --- | --- | --- | --- | --- |
| Species name: | Sample size (n) | Location | Std. Length Range (mm) | Std. Length Avg. | Std. Length S.D. | Gut Length Range (mm) | Gut Length Avg. | Gut Length S.D. | Relative Gut Length Avg. |
| *Cyprinodon variegatus*<br>Generalist | 5 | Crescent Pond, San Salvador Island, Bahamas | 30-38 | 32.6 | 3.13 | 66-94 | 75.6 | 11.61 | 2.32 |
| *Cyprinodon brontotheroides* | 5 | Crescent Pond, San Salvador Island, Bahamas | 26-35 | 30.4 | 4.34 | 49-79 | 64.8 | 12.17 | 2.13 |
| *Cyprinodon desquamator* | 5 | Crescent Pond, San Salvador Island, Bahamas | 28-34 | 30.6 | 2.61 | 49-84 | 65.8 | 14.02 | 2.15 |
| *Cyprinodon variegatus*<br>Generalist | 5 | Osprey Lake, San Salvador Island, Bahamas | 24-38 | 30.6 | 5.18 | 28-95 | 55.4 | 24.64 | 1.81 |
| *Cyprinodon brontotheroides* | 5 | Osprey Lake, San Salvador Island, Bahamas | 26-32 | 28 | 2.10 | 34-66 | 46.2 | 12.70 | 1.65 |
| *Cyprinodon desquamator* | 5 | Osprey Lake, San Salvador Island, Bahamas | 27-31 | 28.8 | 1.48 | 46-56 | 51.4 | 4.22 | 1.78 |
| ***Cyprinodon variegatus***<br>**Generalist** | 2 | Fort Fisher,<br>North<br>Carolina,<br>USA | 33-35 | 34 | 1.14 | 67-72 | 69.5 | 3.54 | 2.04 |
| ***Cyprinodon laciniatus***<br>**Generalist** | 4 | Lake<br>Cunningham,<br>New<br>Providence<br>Island,<br>Bahamas | 36-48 | 41.5 | 5.00 | 83-101 | 84.75 | 11.50 | 2.04 |
| ***Cualac tessellatus***<br>**Generalist** | 4 | San Luis<br>Potosí,<br>Mexico | 25-32 | 29 | 3.16 | 37-54 | 43.5 | 7.42 | 1.5 |

**Supplemental Table S2:**
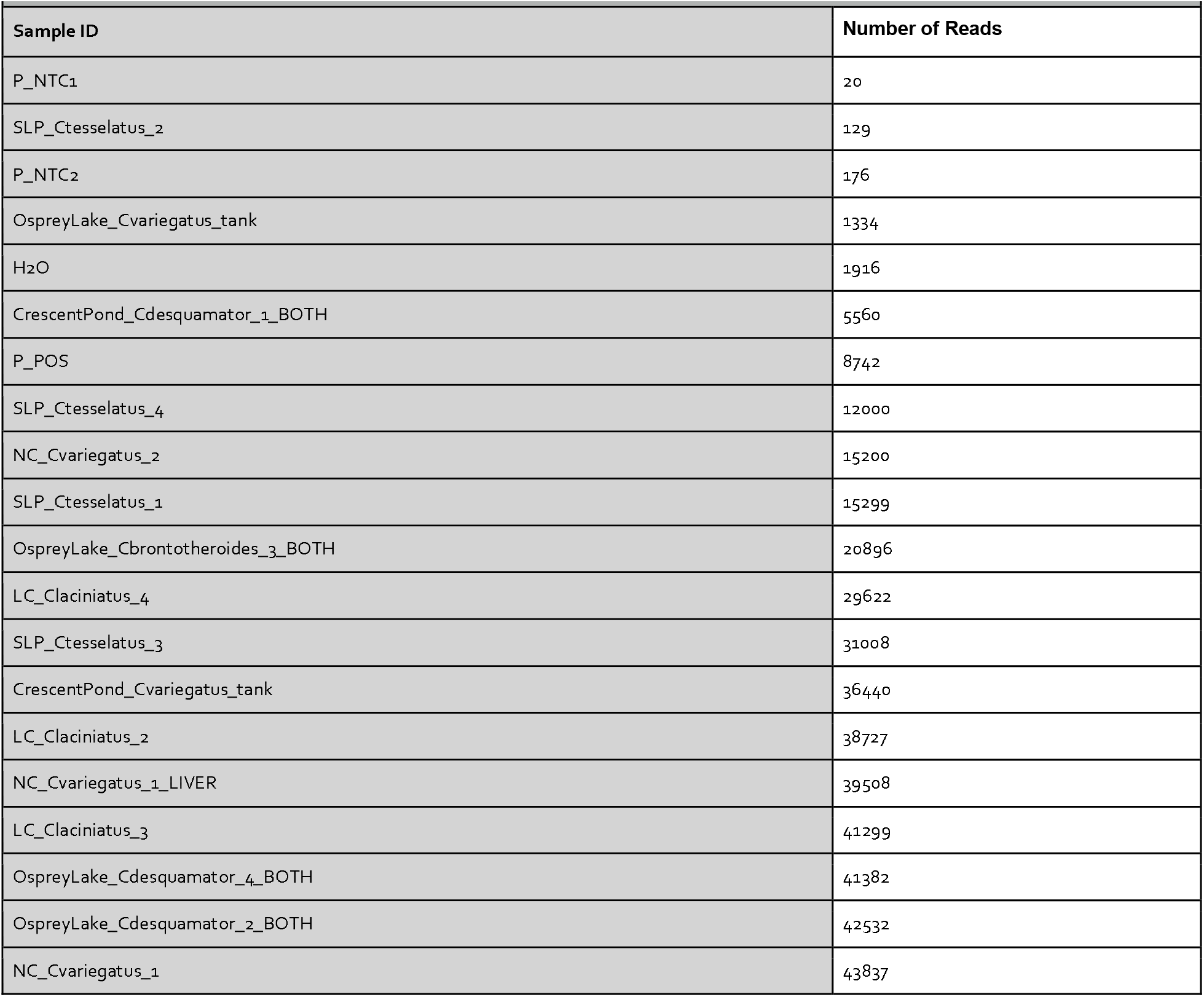
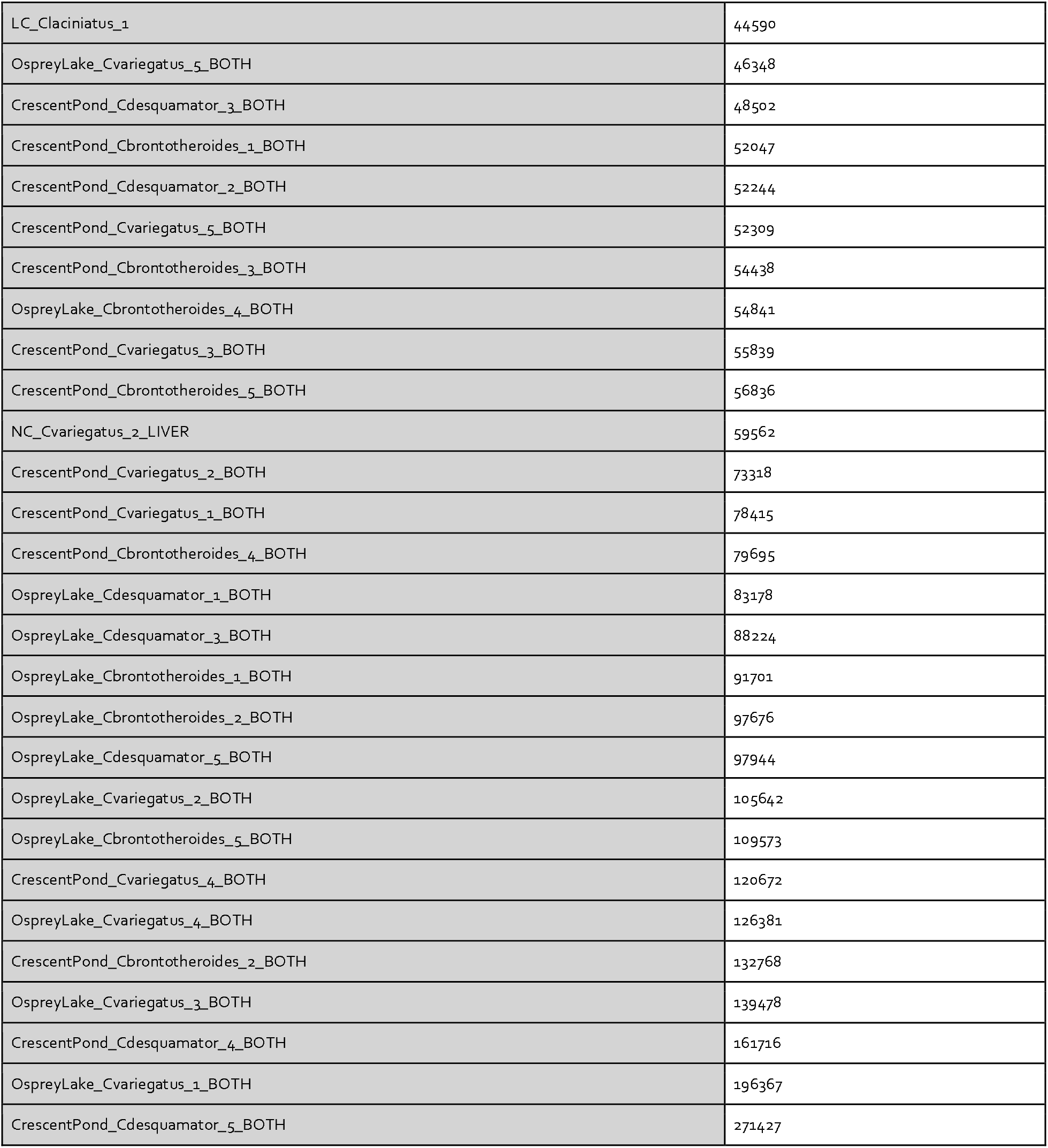
Read Counts (prior to filtering)

| Supplemental Table S2: Read Counts (prior to filtering) |  |
| --- | --- |
| Sample ID | Number of Reads |
| P_NTC1 | 20 |
| SLP_Ctesselatus_2 | 129 |
| P_NTC2 | 176 |
| OspreyLake_Cvariegatus_tank | 1334 |
| H<sub>2</sub>O | 1916 |
| CrescentPond_Cdesquamator_1_BOTH | 5560 |
| P_POS | 8742 |
| SLP_Ctesselatus_4 | 12000 |
| NC_Cvariegatus_2 | 15200 |
| SLP_Ctesselatus_1 | 15299 |
| OspreyLake_Cbrontotheroides_3_BOTH | 20896 |
| LC_Claciniatus_4 | 29622 |
| SLP_Ctesselatus_3 | 31008 |
| CrescentPond_Cvariegatus_tank | 36440 |
| LC_Claciniatus_2 | 38727 |
| NC_Cvariegatus_1_LIVER | 39508 |
| LC_Claciniatus_3 | 41299 |
| OspreyLake_Cdesquamator_4_BOTH | 41382 |
| OspreyLake_Cdesquamator_2_BOTH | 42532 |
| NC_Cvariegatus_1 | 43837 |
| LC_Claciniatus_1 | 44590 |
| OspreyLake_Cvariegatus_5_BOTH | 46348 |
| CrescentPond_Cdesquamator_3_BOTH | 48502 |
| CrescentPond_Cbrontotheroides_1_BOTH | 52047 |
| CrescentPond_Cdesquamator_2_BOTH | 52244 |
| CrescentPond_Cvariegatus_5_BOTH | 52309 |
| CrescentPond_Cbrontotheroides_3_BOTH | 54438 |
| OspreyLake_Cbrontotheroides_4_BOTH | 54841 |
| CrescentPond_Cvariegatus_3_BOTH | 55839 |
| CrescentPond_Cbrontotheroides_5_BOTH | 56836 |
| NC_Cvariegatus_2_LIVER | 59562 |
| CrescentPond_Cvariegatus_2_BOTH | 73318 |
| CrescentPond_Cvariegatus_1_BOTH | 78415 |
| CrescentPond_Cbrontotheroides_4_BOTH | 79695 |
| OspreyLake_Cdesquamator_1_BOTH | 83178 |
| OspreyLake_Cdesquamator_3_BOTH | 88224 |
| OspreyLake_Cbrontotheroides_1_BOTH | 91701 |
| OspreyLake_Cbrontotheroides_2_BOTH | 97676 |
| OspreyLake_Cdesquamator_5_BOTH | 97944 |
| OspreyLake_Cvariegatus_2_BOTH | 105642 |
| OspreyLake_Cbrontotheroides_5_BOTH | 109573 |
| CrescentPond_Cvariegatus_4_BOTH | 120672 |
| OspreyLake_Cvariegatus_4_BOTH | 126381 |
| CrescentPond_Cbrontotheroides_2_BOTH | 132768 |
| OspreyLake_Cvariegatus_3_BOTH | 139478 |
| CrescentPond_Cdesquamator_4_BOTH | 161716 |
| OspreyLake_Cvariegatus_1_BOTH | 196367 |
| CrescentPond_Cdesquamator_5_BOTH | 271427 |

